# Asialoglycoprotein receptor 1 is a novel PCSK9-independent ligand of liver LDLR that is shed by Furin

**DOI:** 10.1101/2021.04.11.439364

**Authors:** Delia Susan-Resiga, Emmanuelle Girard, Rachid Essalmani, Anna Roubtsova, Jadwiga Marcinkiewicz, Rabeb M. Derbali, Alexandra Evagelidis, Jae H. Byun, Paul F. Lebeau, Richard C. Austin, Nabil G. Seidah

## Abstract

The hepatic carbohydrate-recognizing asialoglycoprotein receptor (ASGR1) mediates the endocytosis/lysosomal degradation of desialylated glycoproteins following binding to terminal galactose/N-acetylgalactosamine. Human heterozygote-carriers of *ASGR1-*deletions exhibited ∼34% lower risk of coronary artery disease and ∼10-14% non-HDL-cholesterol reduction. Since PCSK9 is a major degrader of LDLR, the regulation of LDLR and/or PCSK9 by ASGR1 was studied. Thus, we investigated the role of endogenous/overexpressed ASGR1 on LDLR degradation and functionality by Western-blot and immunofluorescence in HepG2 naïve and HepG2-PCSK9-knockout cells. ASGR1, like PCSK9, targets LDLR and both interact with/enhance the degradation of the receptor independently. Such lack of cooperativity between PCSK9 and ASGR1 on LDLR expression was confirmed in livers of wild-type (WT) *versus Pcsk9*^*-/-*^ mice. ASGR1-knockdown in HepG2 naïve cells significantly increased total (∼1.2-fold) and cell-surface (∼4-fold) LDLR protein. In HepG2-PCSK9-knockout cells ASGR1-silencing led to ∼2-fold higher levels of LDLR protein and DiI-LDL uptake, associated with ∼9-fold increased cell-surface LDLR. Overexpression of WT-ASGR1/2 reduced primarily the immature non-O-glycosylated LDLR (∼110 kDa), whereas the triple Gln^240^/Trp^244^/Glu^253^ Ala-mutant (loss of carbohydrate-binding) reduced the mature form of the LDLR (∼150 kDa), suggesting that ASGR1 binds the LDLR in sugar-dependent and -independent fashion. Furin sheds ASGR1 at **<u>R</u>**KM**<u>K</u>**^103^↓ into a secreted form, likely resulting in a loss-of-function on LDLR. LDLR is the first example of a liver-receptor ligand of ASGR1. Additionally, we demonstrate that lack of ASGR1 enhances LDLR levels and DiI-LDL incorporation, independently of PCSK9. Overall, silencing of ASGR1 and PCSK9 may lead to higher LDL-uptake by hepatocytes, thereby providing a novel approach to further reduce LDL-cholesterol.

## INTRODUCTION

Elevated levels of plasma low-density lipoprotein cholesterol (LDLc) and inflammation are the leading modifiable risk factors that drive the development and progression of atherosclerosis, the underlying cause of cardiovascular disease (CVD) (1). The removal of plasma LDLc is primarily mediated by the LDL receptor (LDLR) located on the surface of hepatocytes. The proprotein convertase subtilisin/kexin type 9 (PCSK9) is highly expressed in the liver (2) and, upon its secretion into plasma, it binds to and enhances the degradation of the LDLR in a non-enzymatic fashion (3-6), thereby increasing circulating LDLc. Unlike gain-of-function (GOF) PCSK9 variants (7), loss-of-function (LOF) variants enhance LDLR levels and promote LDLc clearance (4,8). The development of inhibitory human monoclonal antibodies against PCSK9 represents a powerful treatment strategy for the management of CVD, which has been shown to reduce the risk of cardiovascular events in clinical populations (9). Despite these hallmark studies and their utility for lipid lowering to prevent CVD, other surface proteins on hepatocytes that bind to and modulate their interaction with PCSK9 and/or LDLR as well as LDLc uptake have not been thoroughly characterized (6).

The asialoglycoprotein receptor (ASGR) (10) is a hepatocyte type-II transmembrane heterodimeric glycoprotein highly conserved among mammals. It consists of two subunits ASGR1 (major) and ASGR2 (minor) and plays a critical role in serum glycoprotein homeostasis by mediating the endocytosis and lysosomal degradation of desialylated glycoproteins with exposed terminal galactose or N-acetylgalactosamine residues (11). ASGRs are highly expressed in hepatocytes (10). Recent studies showed that heterozygous carriers of the early termination LOF del12 mutation in the ASGR1 gene (1 in 120 persons in the Icelandic study) and another early termination heterozygote LOF ASGR1 mutant (W158X) had lower plasma levels of non-HDL cholesterol than non-carriers (12). Additionally, the del12 mutation was associated with a significant 34% lower risk of CVD. Comparison of plasma LDLc decrease in LOF PCSK9 R46L (−17 mg/dL; ∼30% reduction in CVD risk) with that in LOF ASGR1 del12 mutation (−11 mg/dL) illustrates that reduction in CVD risk observed in this ASGR1 mutant (∼34%) was ∼2-fold greater than would have been predicted (i.e., ∼18%) from the associated modest reduction of LDLc (12). This suggests that the atheroprotective effects of ASGR1 del12 go beyond the lowering of serum LDLc.

Strategies aimed at inhibiting ASGR1 expression/activity are being explored as another approach to lowering plasma LDLc and CVD risk (11). Despite the importance of PCSK9 and ASGR1 in lipid lowering and reduced CVD risk, there are currently no studies to suggest that ASGR1 and PSCK9 act in concert to impact lipid metabolism and/or atherosclerosis by modulating the binding/uptake of LDLc *via* the LDLR. The purpose of the present study was to investigate the possible regulation of the LDLR by ASGR1 and the role of PCSK9 in this process.

## RESULTS

### Expression of ASGR1 and ASGR2 during development and in adult mice, and in human hepatocyte cell lines

Human ASGR is composed of a major subunit (ASGR1) and a minor subunit (ASGR2) (13). Both subunits are type II, single-pass transmembrane proteins. In ASGR1, residues Gln_240_, Trp_244_ and Glu_253_ in the carbohydrate recognition domain (CRD) are critical for binding exposed terminal galactose or N-acetylgalactosamine residues (https://www.uniprot.org/uniprot/P07306) (Fig. 1A). In mouse, whole body *in situ* hybridization histochemistry of ASGR1 and ASGR2 mRNA expression during development and in the adult revealed that both transcripts are mainly expressed in liver starting at embryonic days 17 and 15, respectively, and showed that their expression increases until adulthood (Fig. 1B). A Tabula Muris compendium of single cell RNASeq transcriptome data from 20 mouse organs and tissues (czbiohub.org) (14) revealed that ASGR1 transcripts are exclusively expressed in liver, specifically in hepatocytes (Fig. 1C). We next estimated the relative mRNA expression of ASGR1 and ASGR2 by quantitative RT-PCR in adult mouse liver and in human hepatocyte HepG2, HepG2-PCSK9-KO cells (lacking endogenous PCSK9) (15), and in the immortalized human primary hepatocytes IHH cell line (Fig. 1D) (15). In most cases the mRNA levels of ASGR1 were higher than those of ASGR2, especially in mouse liver.

**Figure 1:**
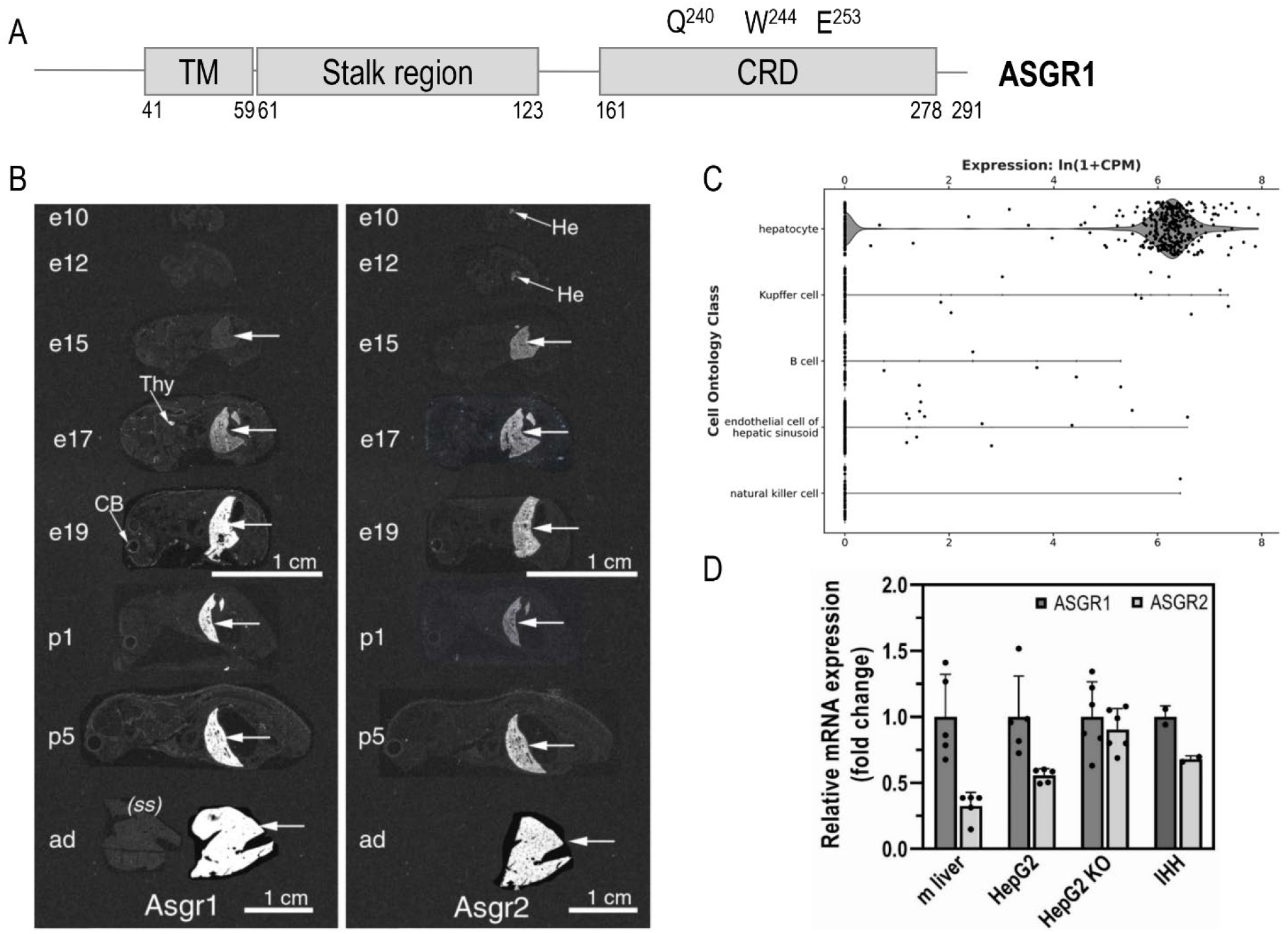
Expression of ASGR1 and ASGR2 in mice and cell lines. (A) Schematic representation of ASGR1, depicting its cytoplasmic N terminus (40 amino acids), the single transmembrane domain (TM) and its exoplasmic C terminus (∼230 aa) that is composed of a stalk region and a C-terminal carbohydrate recognition domain (CRD) with three reported residues important for carbohydrate binding. (B) *In situ* hybridization of ASGR1 and ASGR2 in mice before and after birth. (C) A Tabula Muris Consortium of mouse liver single-cell transcriptomics by RNAseq (czbiohub.org) emphasizes the expression of ASGR1 transcripts in hepatocytes; CPM = counts per million reads mapped. (D) Quantitative RT-PCR of ASGR1 and ASGR2 in mouse (m) liver, HepG2 naïve, HepG2-PCSK9-KO, and IHH cells. For each sample analyzed, the ASGR2 mRNA expression was normalized to the one of ASGR1. Quantifications are averages ± SD.

### PCSK9 reduces LDLR but not ASGR1 levels

Immunofluorescence (IF) staining of endogenous ASGR1 in HepG2-PCSK9-KO cells confirmed its presence at the plasma membrane and revealed its co-localization with the LDLR (-PCSK9; Fig 2A). Circulating PCSK9 is responsible for most of its activity to reduce the levels of cell-surface LDLR in hepatocytes (6). Hence, we addressed the potential regulation of ASGR1 by exogenous purified recombinant PCSK9 (16). We thus compared the known ability of *exogenous* PCSK9 to enhance the degradation of LDLR to its effect on ASGR1 in HepG2 cells by IF (Fig. 2A), and Western blot (WB; Fig. 2B). As expected, PCSK9 reduced the endogenous levels of cell-surface LDLR (−90% by IF) and total LDLR (−40% by WB). In contrast, PCSK9 did not alter the levels of endogenous ASGR1 in both assays. Note the presence of two bands for endogenous ASGR1 (Fig. 2B), likely representing a major mature form (∼48 kDa) that excited the endoplasmic reticulum (ER) and a minor ER-localized immature form (∼42 kDa), as observed in other proteins (16,17).

**Figure 2:**
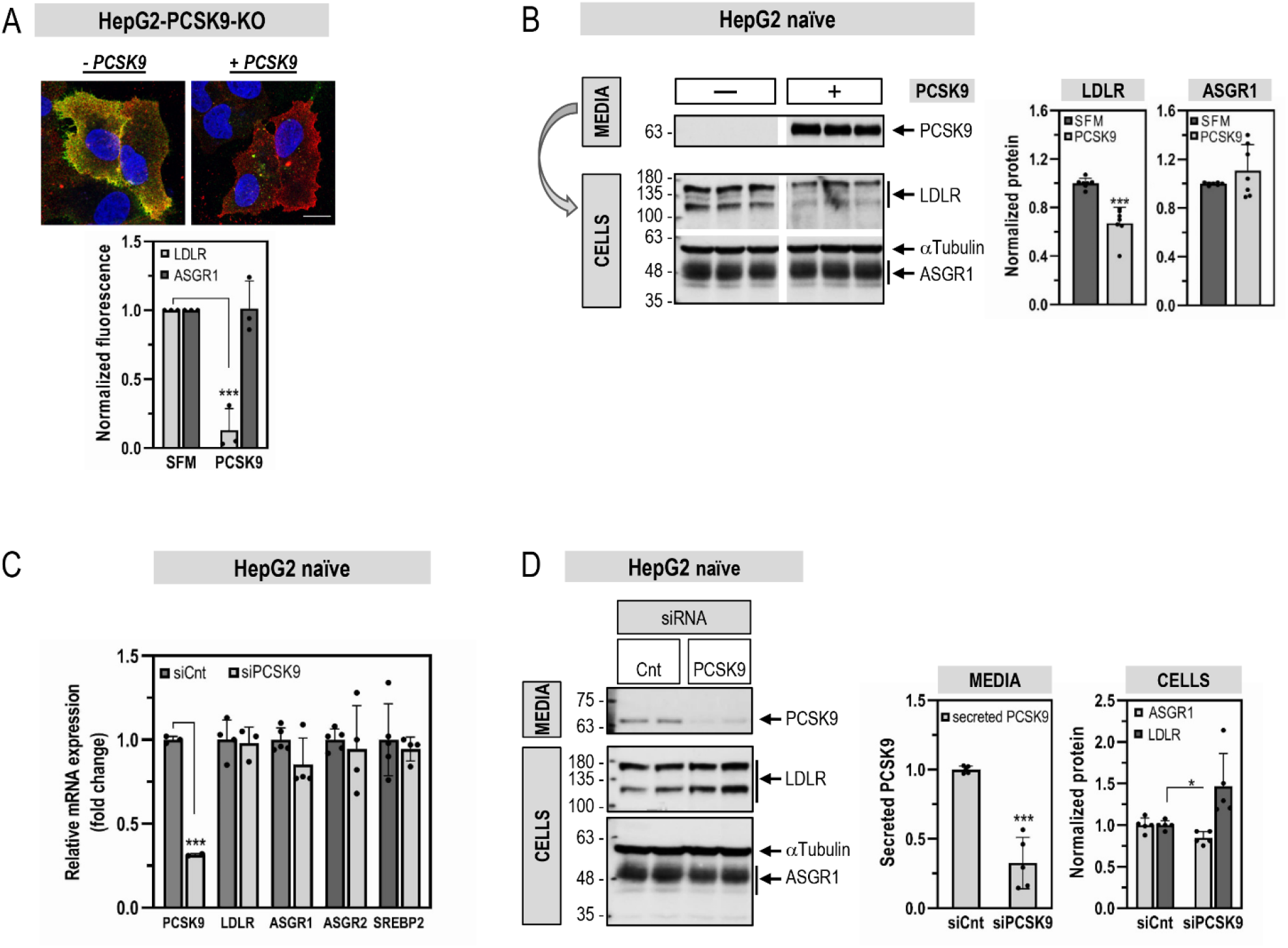
Comparative effects of PSCK9 on cellular LDLR and ASGR1 levels. HepG2-PCSK9-KO cells (A) or HepG2 naïve cells (B) were incubated for 20 hours with media only (-PCSK9; SFM) or media containing purified PCSK9 (3 μg/ml) (+ PCSK9). (A) Immunofluorescence microscopy of plasma membrane LDLR (green signal) and ASGR1 (red signal). The co-localization appears as yellow. Scale bar, 15 µm. (B) WB analysis of cellular LDLR and ASGR1 expression and of PCSK9 in the media. (C, D) HepG2 naïve cells were transfected with non-targeting siRNA (siCnt) or siRNA PCSK9 (siPCSK9) at final concentrations of 10 nM and analyzed 48 hours post-transfection. (C) Quantitative RT-PCR analysis. (D) WB analysis of cellular LDLR and ASGR1 and of secreted PCSK9. Data are representative of at least three independent experiments. Quantifications are averages ± SD. * p<0.05; *** p<0.001 (t-test).

The effect of *endogenous* PCSK9 on LDLR and ASGR1 in HepG2 cells was next investigated following its mRNA knockdown (KD) using a smart pool of 4 siRNAs. This resulted in ∼70% reduction in PCSK9 mRNA and protein levels (Figs. 2C, D). As expected (18,19), lack of PCSK9 did not affect the mRNA levels of LDLR. In addition, no effect was observed on the mRNA levels of ASGR1, ASGR2 and the transcription factor SREBP2 implicated in PCSK9 mRNA regulation (20) (Fig. 2E). At the protein level, PCSK9 KD resulted in ∼1.5-fold increase in LDLR, with no change in ASGR1 (Fig. 2D). We conclude that in HepG2 cells *endogenous* and *exogenous* PCSK9 do not affect ASGR1 mRNA and/or protein levels.

To address the effect of PCSK9 on ASGR1 *in vivo*, we analyzed the levels of ASGR1 in *Pcsk9*-knockout (KO) mice (18), by liver immunohistochemistry (IHC) under non-permeable conditions (21) (Fig. 3A) and WB (Fig. 3B). The data showed that while LDLR protein levels at the cell-surface of hepatocytes (IHC) and in whole tissue (WB) were increased in *Pcsk9*-KO livers, in accordance with (21), those of ASGR1 were not affected. Overall, we conclude that, like in HepG2 cells, *in vivo* endogenous PCSK9 does not affect the levels of ASGR1 in mouse hepatocytes.

**Figure 3.**
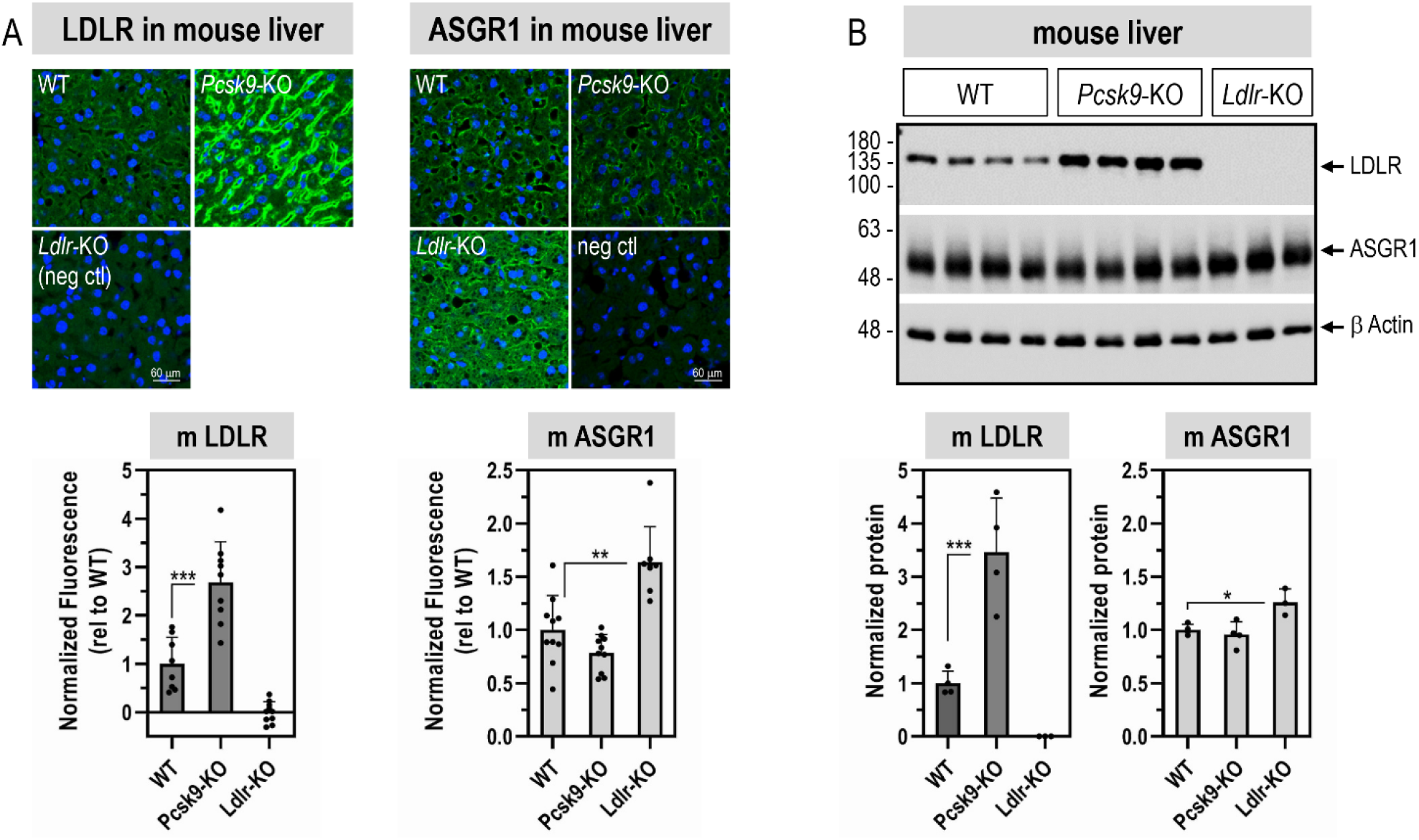
: LDLR and ASGR1 expression levels in livers from wild-type (WT) mice and mice lacking mPCSK9 (*PCSK9*-KO) or mLDLR (*Ldlr*-KO), respectively. (A) LDLR and ASGR1 staining by immunohistochemistry (green signal) in mouse liver sections. (B) WB analysis of total ASGR1 and LDLR proteins in mouse livers. Data are representative of 9-10 mice (A) and 3-4 mice (B) and quantifications are averages ± SD. * p<0.05; ** p<0.01; *** p<0.001 (t-test).

To analyze the impact of the lack of LDLR on ASGR1 protein, we now show that in the liver of *Ldlr*-KO mice both ASGR1 cell-surface immunoreactivity (Fig. 3A) and total ASGR1 protein by WB (Fig. 3B) were significantly increased by ∼1.6 and ∼1.3 fold, respectively. This suggests that ASGR1 levels could be regulated by those of the LDLR, as are circulating PCSK9 levels (6).

### ASGR1 regulates LDLR levels and functionality independently of PCSK9

To examine the impact of ASGR1 on the LDLR, KD of ASGR1 (−75%) was performed in naïve HepG2 and HepG2-PCSK9-KO cells (Figs. 4, 5). We first demonstrated that KD of ASGR1 in naïve HepG2 cells did not affect the levels of intracellular and secreted PCSK9 (Fig. 4). In contrast, ASGR1 KD resulted in a significant increase in total (WB) and cell-surface (IF) LDLR in both cell lines. Specifically, we estimated a total LDLR increase of ∼1.3-fold (Fig. 5A) and ∼2-fold (Fig. 5B) in each cell line, respectively. The corresponding cell-surface increase in LDLR was at least ∼4-fold in naïve HepG2 cells (Figs. 5C, D) and ∼9-fold in HepG2-PCSK9-KO cells (Fig. 5E). Transcript analysis by quantitative RT-PCR revealed that KD of ASGR1 in HepG2 cells did not modify the mRNA levels of LDLR and SREBP2 (Fig. 5F). This suggests that endogenous ASGR1 reduces the total levels of LDLR protein, especially those at the cell-surface, leading to the hypothesis that LDLR is a target of ASGR1, independent of PCSK9.

**Figure 4:**
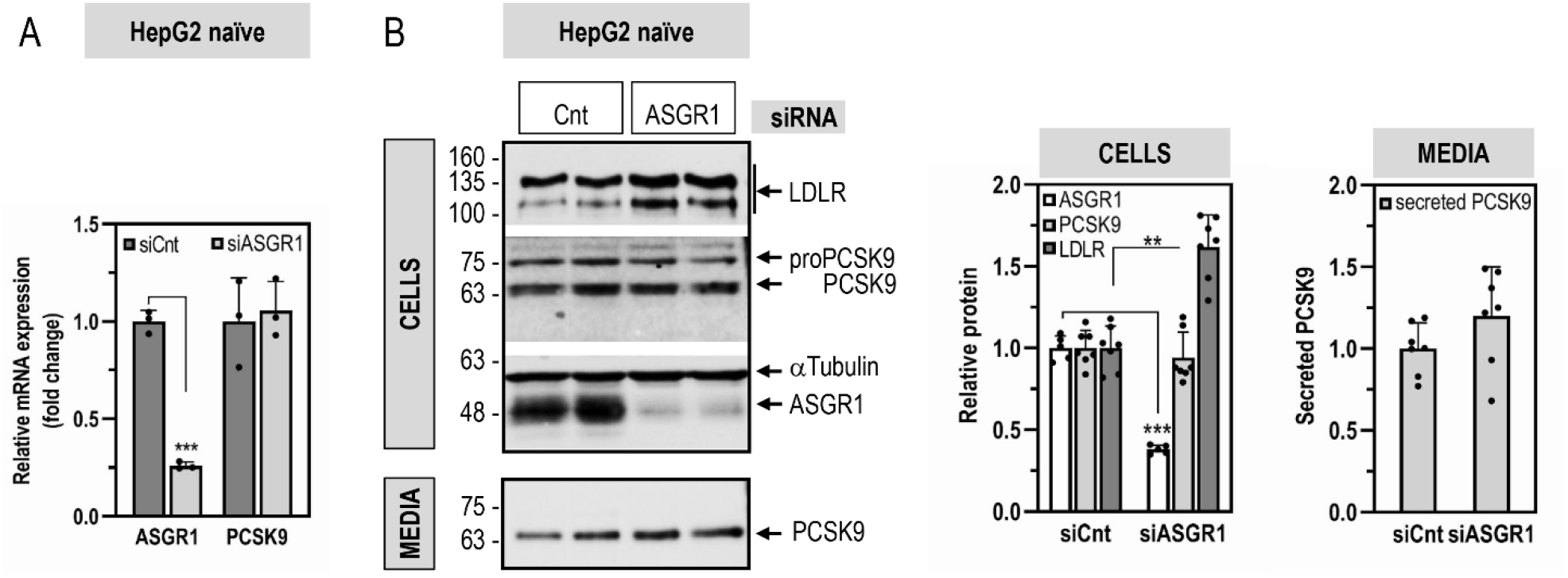
ASGR1 does not affect PCSK9 expression and secretion. HepG2 naïve cells were transfected for 48 hours with non-targeting siRNA (siCnt) or siRNA ASGR1 (siASGR1) at final concentrations of 20 nM and analyzed by quantitative RT-PCR (A), or WB for cellular LDLR, PCSK9, and ASGR1 and for secreted PCSK9 (B). Data are representative of at least three independent experiments. Quantifications are averages ± SD. ** p<0.01; *** p<0.001 (t-test).

**Figure 5:**
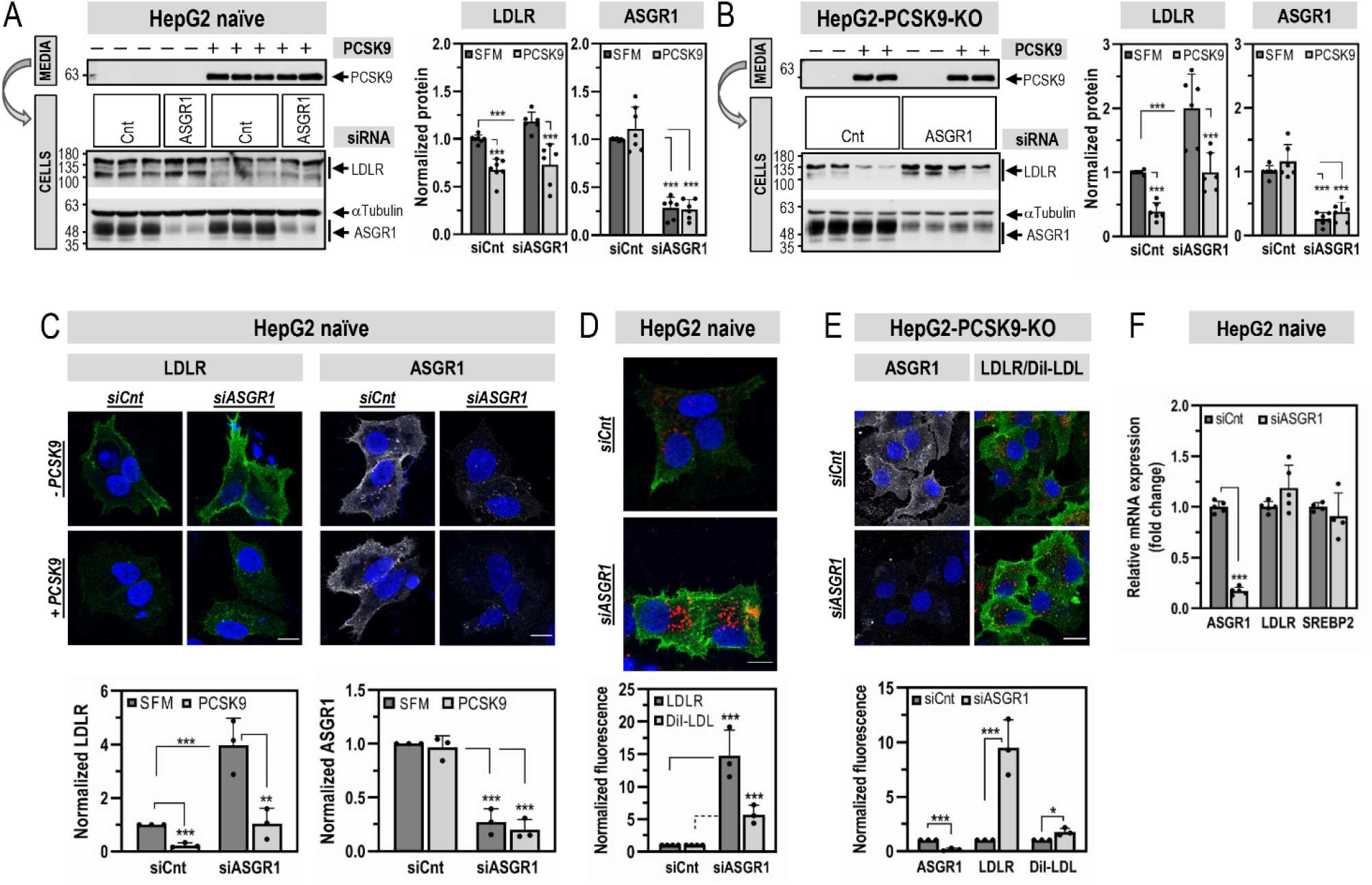
ASGR1 regulates LDLR levels and functionality independently of PCSK9. HepG2 naïve cells (A, C, D) or HepG2-PCSK9-KO cells (B, E) were transfected for 48 hours with non-targeting siRNA (siCnt) or siRNA ASGR1 (siASGR1) at final concentrations of 20 nM. Cells were then incubated for the last 20 hours with media only (-PCSK9; SFM) or media containing purified PCSK9 (3 μg/ml) (+ PCSK9) (A-C). (A-B) Total protein levels of LDLR and ASGR1 were assessed by WB. (C) Immunofluorescence microscopy of LDLR (green signal) and ASGR1 (white signal) at the plasma membrane. (D, E) Immunofluorescence microscopy of plasma membrane LDLR (green signal) and (E) plasma membrane ASGR1 (white signal). LDLR functionality was assessed by DiI-LDL (5 μg/ml) internalization for 2 hours at 37°C before fixation (red signal). Scale bar, 15 μm. (F) Quantitative RT-PCR analysis was performed on HepG2 naïve cells transfected for 48 hours with non-targeting siRNA (siCnt) or siRNA ASGR1 (siASGR1) at final concentrations of 20 nM. Data are representative of at least three independent experiments. Quantifications are averages ± SD. * p<0.05; ** p<0.01; *** p<0.001 (t-test).

We next investigated whether ASGR1 affects the ability of PCSK9 to enhance the degradation of the LDLR. In both cell lines KD of ASGR1 had no significant impact on the activity of exogenous PCSK9 on the LDLR (Figs. 5A, B, C). Thus, we can conclude that independently PCSK9 and ASGR1 can reduce the levels of LDLR protein.

To probe the functionality of the LDLR upon KD of ASGR1, we incubated naïve HepG2 (Fig. 5D) and HepG2-PCSK9-KO cells (Fig. 5E) with fluorescent DiI-LDL and estimated its uptake by IF. The data show that the KD of ASGR1 resulted in at least ∼2-fold increase in DiI-LDL uptake mirroring the increase in LDLR protein levels. Thus, endogenous ASGR1 negatively regulates protein levels of LDLR and its functionality independently of PCSK9.

Since ASGR1 can act as a homodimer or heterodimer with ASGR2 (10), we next examined the effect of overexpression of ASGR1/2 in HepG2-PCSK9-KO cells. Co-expression of ASGR1 and ASGR2 at a 1:1 ratio resulted in a significant decrease in the total levels of both endogenous (−30%) and overexpressed (−55%) LDLR (WB; Fig. 6A). Similarly, overexpressed ASGR1/2 reduced the levels of endogenous and overexpressed cell-surface LDLR (−75%) (IF; Fig. 6B). Notably, HepG2-PCSK9-KO cells endogenously express ASGR1 and ASGR2 (Fig. 1D). Thus, their individual overexpression would be expected to result in homodimers and/or heterodimers with the endogenous ASGRs, consequently affecting LDLR levels. This was evident in IF experiments where analyses were focused on transfected cells only (Fig. 6B), and in WB analyses with mAb-V5 of overexpressed LDLR-V5 (Fig. 6A; right panel). In all the above, overexpression of either ASGR1, or ASGR2 or their combination gave similar results. Altogether, KD and overexpression analyses in HepG2-PCSK9-KO cells demonstrated that, independent of PCSK9, ASGR1 negatively regulates LDLR levels and its functionality.

**Figure 6:**
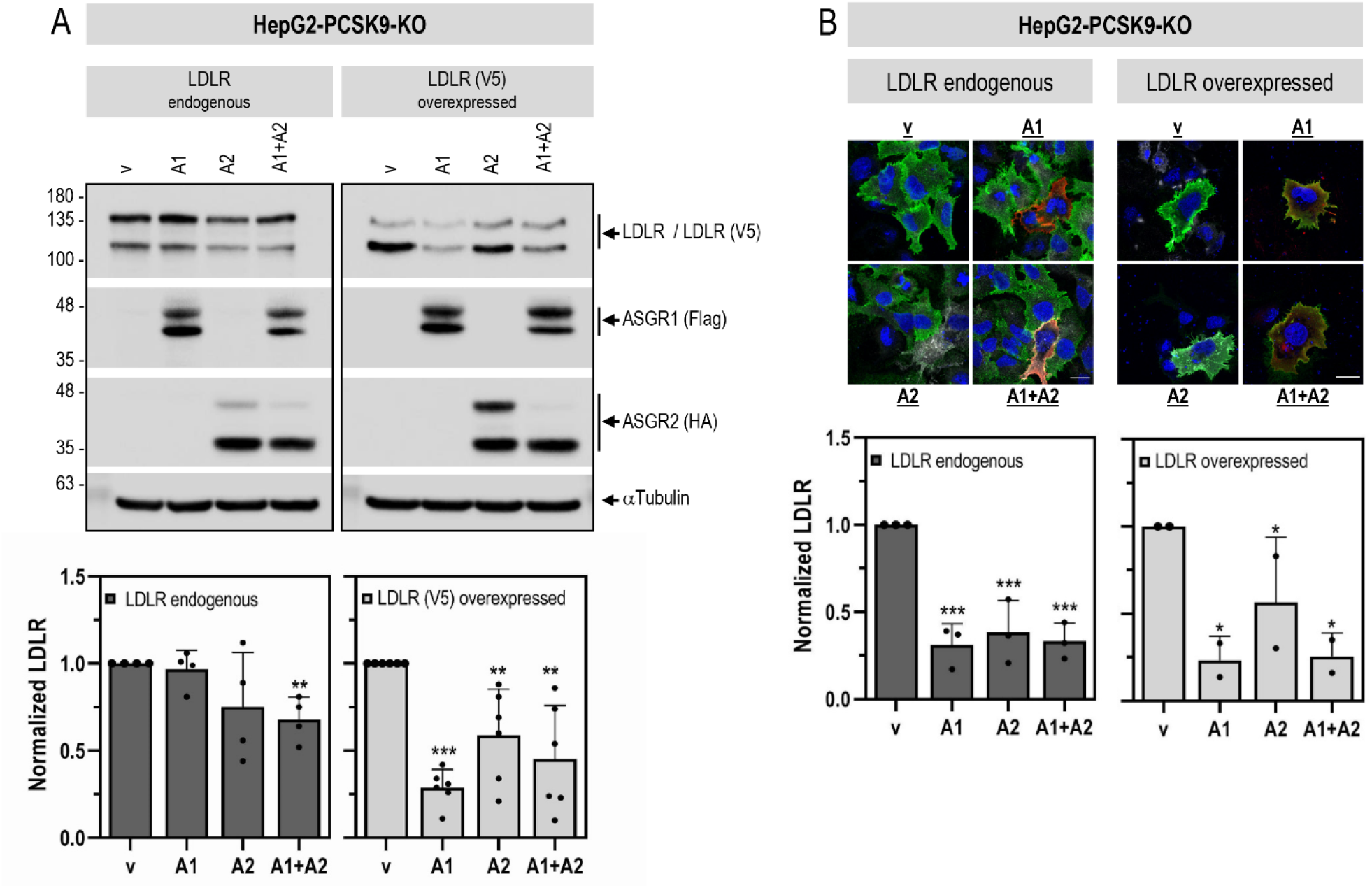
Overexpressed ASGR1 and ASGR2 reduce LDLR levels independently of PCSK9. HepG2-PCSK9-KO cells were transfected with either Flag-tagged ASGR1 (A1), HA-tagged ASGR2 (A2) or their combination (A1+A2), or control empty vector (v), alone or in combination with LDLR(V5). (A) WB analyses and quantification of cellular LDLR (LDLR endogenous: LDLR antibody; LDLR (V5) overexpressed: V5 antibody), ASGR1 (Flag-HRP) and ASGR2 (HA-HRP). (B) Immunofluorescence microscopy of endogenous or overexpressed cell-surface LDLR (green signal). Quantification was performed only in cells transfected with ASGR1 (red signal) or/and ASGR2 (white signal). Scale bar, 15 μm. Data are representative of at least two independent experiments. Quantifications are averages ± SD. * p<0.1; ** p<0.01; *** p<0.001 (t-test) are relative to the control vector condition.

To dissect the ASGR1-induced degradation of the LDLR in the presence or absence of ASGR2, we selected HEK293 cells that do not express ASGR1/2 endogenously. The data showed that overexpression of ASGR1 or ASGR2 alone or together (at a 1:1 ratio) led to the same ∼30% decrease in total LDLR-V5 levels (Fig. 7). These results suggest that under overexpression conditions both ASGR1 and ASGR2 can degrade the LDLR, likely as homo- or hetero-oligomers (10). Hence, to maintain closer to *in vivo* conditions where both ASGR1/2 are present, we chose to use co-expressed ASGR1/2 in all subsequent experiments.

**Figure 7:**
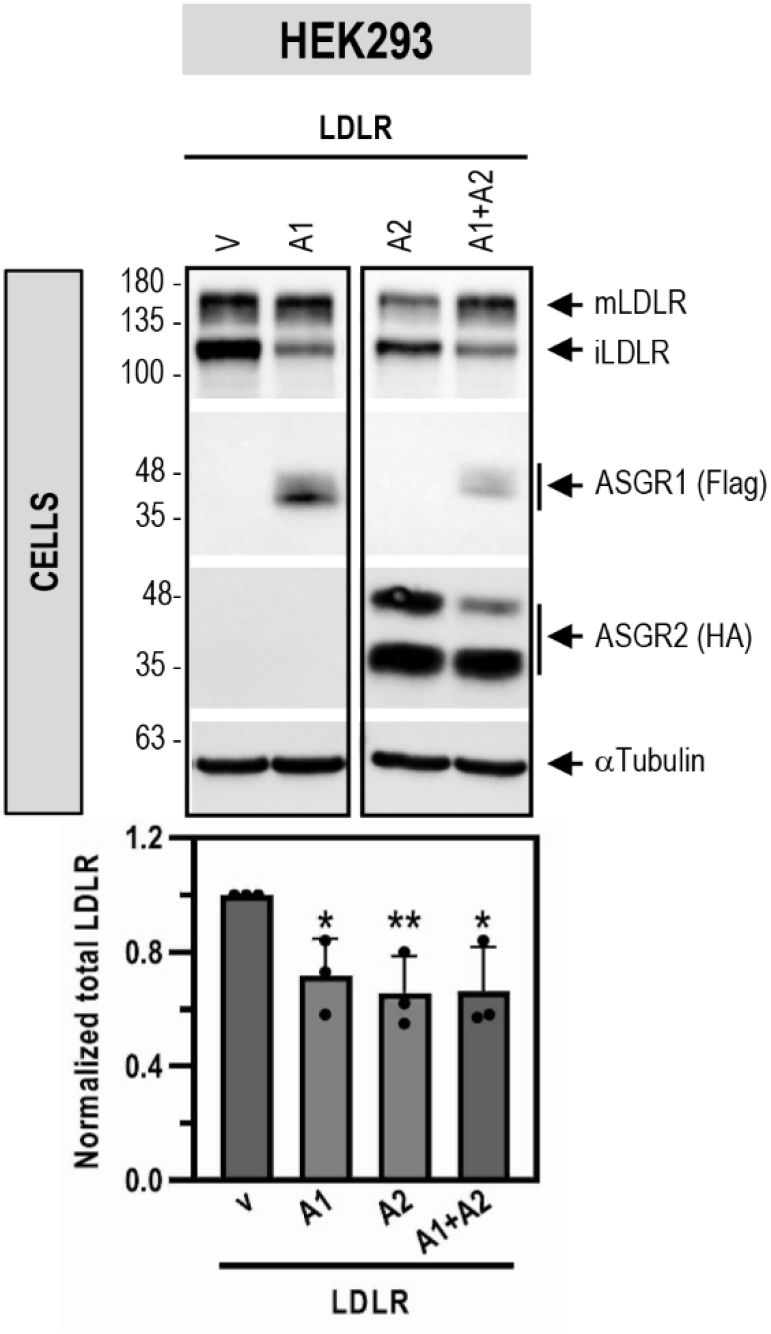
ASGR1 and ASGR2 degrade LDLR under overexpression conditions in HEK293 cells. HEK293 cells were co-transfected with V5-tagged LDLR and either Flag-tagged ASGR1 (A1), HA-tagged ASGR2 (A2) or their combination (A1+A2), or control empty vector (v). WB analyses and quantification of cellular LDLR(V5) (V5 antibody), ASGR1 (Flag-HRP) and ASGR2 (HA-HRP) are shown. Data are representative of three independent experiments. Quantifications are averages ± SD. * p<0.1; ** p<0.01 (t-test) are relative to the control vector condition.

### ASGR1 can bind LDLR and enhance its degradation by carbohydrate-dependent and independent mechanisms

Co-immunoprecipitation analyses were used to probe the possible interaction between endogenous ASGR1 and LDLR in HepG2-PCSK9-KO cells (Fig. 8A). Thus, immunoprecipitation of cell lysates under non-denaturing conditions with an ASGR1 antibody, followed by separation of the immune complex by SDS-PAGE and WB revealed that human ASGR1 co-immunoprecipitated (co-IP) with endogenous LDLR (mostly present as mature ∼150 kDa mLDLR), but not with CD36 or insulin receptor (IR, detected with an antibody to the β-subunit), two other endogenous membrane-bound N-glycosylated proteins (Fig. 8A). This suggests that endogenous ASGR1 and LDLR interact in a specific manner and form a complex independent of PCSK9. This observation was also confirmed in *vivo*, in mouse liver from wild type (WT) and *Pcsk9*-KO mice (Fig. 8B).

**Figure 8:**
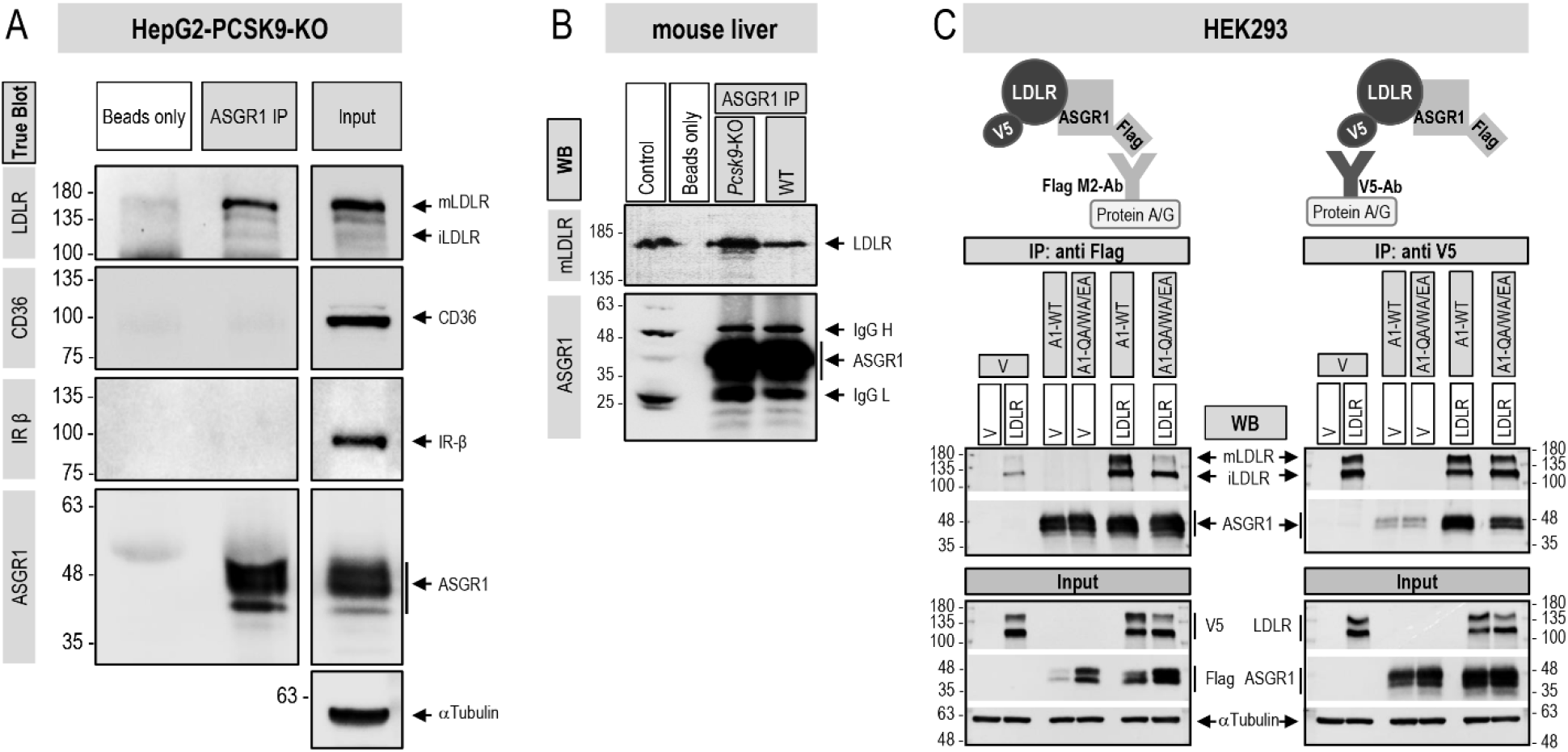
LDLR and ASGR1 form a complex that is not limited to lectin binding. (A) Co-immunoprecipitation of endogenous LDLR and ASGR1 in HepG2-PCSK9-KO cells. Pull-downs with ASGR1 antibody were analyzed by TrueBlot for ASGR1 and LDLR, and for CD36 and IR (shown as IR-β) as negative controls for membrane-bound receptors. To “Beads only” negative control addition of ASGR1 antibody was omitted. “Input” represents 10% of the original material subjected to co-immunoprecipitation. (B) Co-immunoprecipitation from liver extracts of WT and *PCSK9*-KO mice. Pull-downs with ASGR1 antibody were analyzed by WB for LDLR and ASGR1. To “Beads only” negative control addition of ASGR1 antibody was omitted. “Control” (IgG migration control) consists of WT mouse liver lysate (30 µg) plus ASGR1 antibody. (C) Co-immunoprecipitation from HEK293 cells transfected with either vector (v), V5-tagged LDLR or Flag-tagged ASGR1, WT or mutant Q240A/W244A/E253A (QA/WA/EA), or a combination of ASGR1 and LDLR. Pull-downs with Flag M2 antibody (left panel) or V5 antibody (right panel) were analyzed by WB for LDLR and ASGR1. “Input” represents 10% of the original material subjected to co-immunoprecipitation. Data are representative of two independent experiments.

ASGR1 is a lectin-binding protein and Gln_240_, Trp_244_ and Glu_253_ in its CRD are critical for carbohydrate binding. Thus, we next probed the interaction of WT ASGR1 or its triple Ala-mutant (QA/WA/EA) with co-expressed LDLR-V5 in HEK293 cells. Immunoprecipitation with Flag antibody (ASGR1-Flag) and WB with antibodies for ASGR1 or LDLR (Fig. 8C; left panel), revealed that WT ASGR1 binds to both immature iLDLR (non-O-glycosylated; ∼110 kDa) and mature mLDLR (N- and O-glycosylated; ∼150 kDa) (22). In contrast, the triple Ala-mutant that lost its carbohydrate-binding capacity, primarily binds the non-O-glycosylated iLDLR (Fig. 8C; left panel). Binding of the LDLR to ASGR1 was further confirmed in a reciprocal binding assay where the LDLR-V5 is immunoprecipitated first with a mAb-V5 and the immune complex is then separated by SDS-PAGE and revealed by WB to ASGR1 (Fig. 8C; right panel).

We next investigated whether the enhanced degradation of functional LDLR requires the intact carbohydrate-binding motif of ASGR1. Thus, the levels of cell-surface LDLR and DiI-LDL uptake were estimated in HEK293 cells expressing either LDLR alone or LDLR and ASGR2 together with either WT ASGR1 or its triple Ala-mutant (QA/WA/EA). In non-permeabilized cells, IF showed that both cell-surface LDLR and DiI-LDL uptake are significantly reduced in the presence of WT ASGR1, but not of its QA/WA/EA triple Ala-mutant (Fig. 9A). Thus, functional cell-surface LDLR levels are reduced only by WT ASGR1 and require its critical carbohydrate-binding residues. We extended these data by biotinylating cell-surface proteins of the above HEK293 cells and analyzed the biotinylated proteins by WB. The data showed that the ratio of cell-surface [iLDLR] / [total LDLR] is significantly reduced (∼40%) by WT ASGR1 but not by the triple mutant (Fig. 9B; lower left panel). Furthermore, we noticed that the cell-surface levels of the triple-Ala mutant of ASGR1 are ∼60% (∼1.6 fold) higher than the WT (Fig. 9B; top left panel), mirroring its ∼70% (∼1.7 fold) increased total cellular levels (Fig. 9B; right panel), and suggesting that even at higher levels, this mutant that binds iLDLR (Fig. 8C; left panel) does not efficiently reduce LDLR levels (Figs. 9A, B). Collectively, the data suggest that the integrity of ASGR1 CRD regulates its ability to reduce the levels of cell-surface iLDLR.

**Figure 9:**
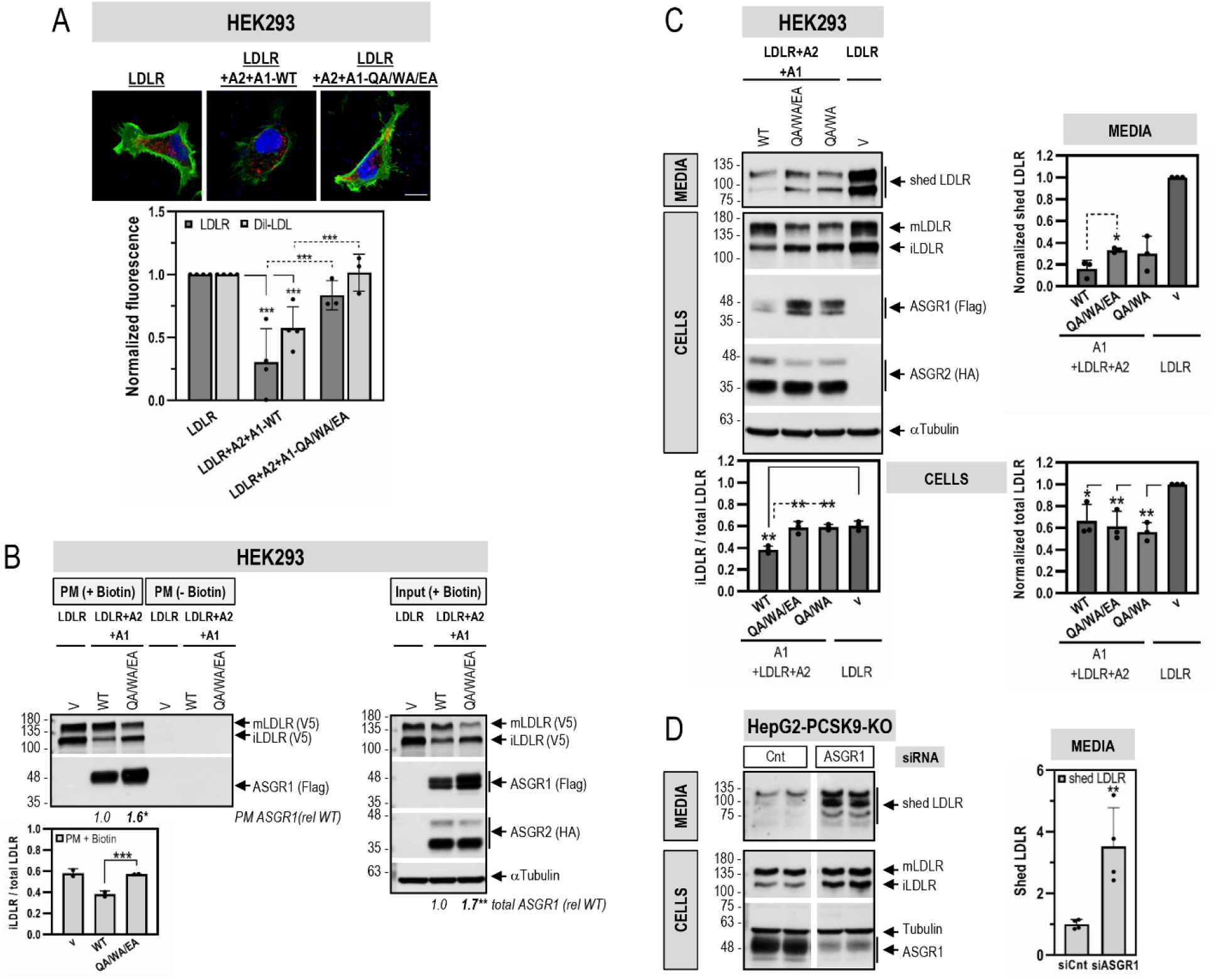
The lectin binding domain in ASGR1 is important for the ASGR1-mediated degradation of the functional, mainly non-O-glycosylated, LDLR at the plasma membrane. HEK293 cells were transfected with V5-tagged LDLR alone or in combination with HA-tagged ASGR2 and Flag-tagged ASGR1, WT or its carbohydrate-binding mutants Q240A/W244A/E253A (QA/WA/EA) (A-C) or Q240A/W244A (QA/WA) (C). (A) Immunofluorescence microscopy of plasma membrane overexpressed LDLR (green signal) and DiI-LDL uptake (red signal) for 2 hours at 37°C before fixation. Non-transfected cells (v) expressed ∼100-fold less LDLR than transfected ones. Scale bar, 15 μm. (B) Surface protein biotinylation: cells incubated with Biotin (+ Biotin) or PBS (-Biotin) were lysed and plasma membrane (PM) proteins were pulled-down with streptavidin and analyzed by WB for LDLR (V5-HRP) and ASGR1 (Flag-HRP) (left panel). One eighth (12.5%; 30 μg) of the (+Biotin) cell lysate before streptavidin pull-down (input) was analyzed by WB (right panel). (C) WB analyses of total cellular LDLR (mature mLDLR, and immature iLDLR), ASGR1 (Flag-HRP) and ASGR2 (HA-HRP), and of shed LDLR in the media. (D) HepG2-PCSK9-KO cells were transfected for 48 hours with 20 nM non-targeting (Cnt) siRNA or siRNA ASGR1. WB analyses of total cellular LDLR (mLDLR and iLDLR), ASGR1 and of shed LDLR in the media are shown. Note that 13% of total media and 8% of total lysates are loaded on the gel and that the exposure times are 15 min and 3 min, respectively. Data are representative of at least two independent experiments. Quantifications are averages ± SD. * p<0.05; ** p<0.01; *** p<0.001 (t-test).

In contrast, WB analyses of the total levels of LDLR in these cells showed that as for WT, the triple QA/WA/EA and double QA/WA mutants of ASGR1 can also significantly reduce total LDLR levels (∼ 40%) relative to a vector control lacking ASGR1/2 (Fig 9C; lower right panel). As observed at the plasma membrane (Fig. 9B; left panel), in total cell lysate (intracellular and plasma membrane) WT ASGR1 also reduced the levels of iLDLR, which is not seen with the CRD mutants that primarily reduce the levels of mLDLR (Fig 9C; lower left panel). This is evident from the relative ratio of [iLDLR] / [total LDLR], which is ∼40% lower in WT ASGR1 *versus* double or triple mutants and vector control (Fig. 9C; lower left panel). Since some of the LDLR is shed into the medium by undefined matrix metalloproteases (23), we also analyzed the fate of media LDLR. The data showed that ASGR1 also reduces the levels of the shed forms of LDLR by carbohydrate-dependent (WT) and carbohydrate-independent (Ala-mutants) activities (Fig. 9C; top right panel). Similar analyses in HEK293 cells only overexpressing ASGR1 and/or LDLR-V5 (Supplementary Fig. S1) mirrored the above results where both ASGR1 and ASGR2 are co-expressed (Fig. 9C). This is taken as evidence that under overexpression conditions, the ASGR1-mediated activity on LDLR *via* its lectin-binding domain is independent of ASGR2. Indeed, silencing ASGR1 in HepG2-PCSK9-KO cells (that still endogenously express ASGR2, Fig. 1D) significantly increased total cellular LDLR levels, associated with ∼3-fold enhanced shedding of the LDLR (∼10% of total LDLR), especially iLDLR (Fig. 9D).

### ASGR1 specificity for LDLR

The present data are the first to identify the secretory membrane-bound LDLR as a physiological target of ASGR1, as opposed to soluble desialylated glycoproteins, e.g., von Willebrand factor (24), normally thought to be sensitive to hepatocyte-derived ASGR1 regulation (11). However, since we do not know the repertoire of liver glycoproteins that are targeted by ASGR1, we next investigated the PCSK9-independent ability of ASGR1 to enhance the degradation of other membrane-bound liver receptors. Accordingly, we analyzed by WB the endogenous levels of CD36 and IR-β following KD of ASGR1 in HepG2-PCSK9-KO cells. The data show that although KD of ASGR1 increased the levels of LDLR by ∼2-fold, those of CD36 and IR-β were not affected (Fig. 10). Thus, even though like the LDLR, the highly N-glycosylated CD36 is a target of PCSK9 (25), it is not sensitive to ASGR1 in the absence of PCSK9. We conclude that the ability of ASGR1 to enhance the degradation of the LDLR may not apply to all liver-derived secretory membrane-bound receptors.

**Figure 10:**
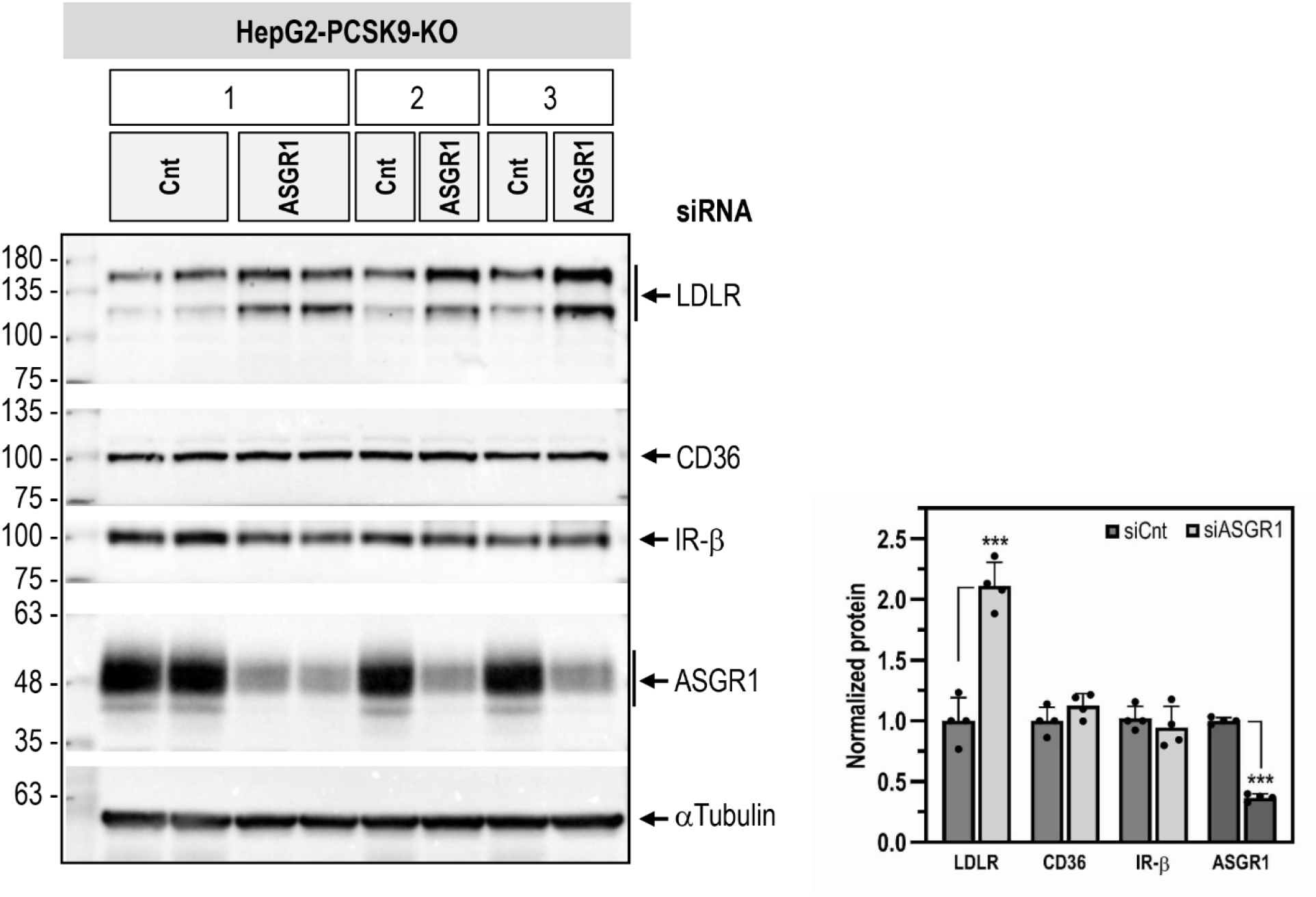
ASGR1 specificity for LDLR. HepG2-PCSK9-KO cells were transfected with 20 nM non-targeting (Cnt) siRNA or siRNA ASGR1 for 48 hours. WB analyses and quantification of total cellular LDLR, ASGR1, CD36 and IR-β from three independent experiments are shown. Quantifications are averages ± SD. *** p<0.001 (t-test).

### Furin sheds ASGR1

Furin, the third member of the proprotein convertase (PC) family (26), has been shown to enhance the shedding of various cell-surface proteins at the general PC-like motif (**<u>R/K</u>**)Xn(**<u>R/K</u>**)↓, where Xn represents 0, 2, 4 or 6 spacer aa (27). These include the type-II membrane-bound CASC4 and GPP130 implicated in cancer/metastasis (28) and the type-I membrane-bound SARS-CoV-2 spike glycoprotein (17). It can also cleave and inactivate PCSK9 in liver hepatocytes (29,30). Scanning the primary sequence of the luminal domain of human ASGR1 revealed the presence of two potential Furin-like cleavage sites, namely **<u>R</u>**GL**<u>R</u>**_74_↓ET and **<u>R</u>**KM**<u>K</u>**_103_**↓**SL. Hence, we addressed the possibility that Furin and/or other PCs could shed ASGR1. Co-expression in HEK293 cells of human ASGR1 with each of the liver-expressed PCs (26) revealed that only Furin can cleave the protein into a soluble ∼28 kDa secreted form (shed ASGR1; Fig. 11A). Whereas Ala-mutation of the **<u>R</u>**GL**<u>R</u>**_74_↓ET site had no effect (Fig. 11B), only Ala-mutation of the underlined basic aa in the **<u>R</u>**KM**<u>K</u>**_103_**↓**SL site abrogated Furin-cleavage and hence shedding of ASGR1 (Fig. 11C). Indeed, insertion of an optimal Furin-like site **<u>RRRR</u>**_103_**↓<u>E</u>**L (17,29) resulted in complete cleavage of mature ∼48 kDa ASGR1 (upper band, likely endoH resistant (17)) by endogenous Furin into a soluble secreted ∼28 kDa sASGR1 (Fig. 11D). A comparable processing pattern was observed in hepatic HepG2-PCSK9-KO cells (Fig. 11E). In conclusion, Furin sheds ASGR1, possibly resulting in a loss-of-function towards LDLR.

**Figure 11:**
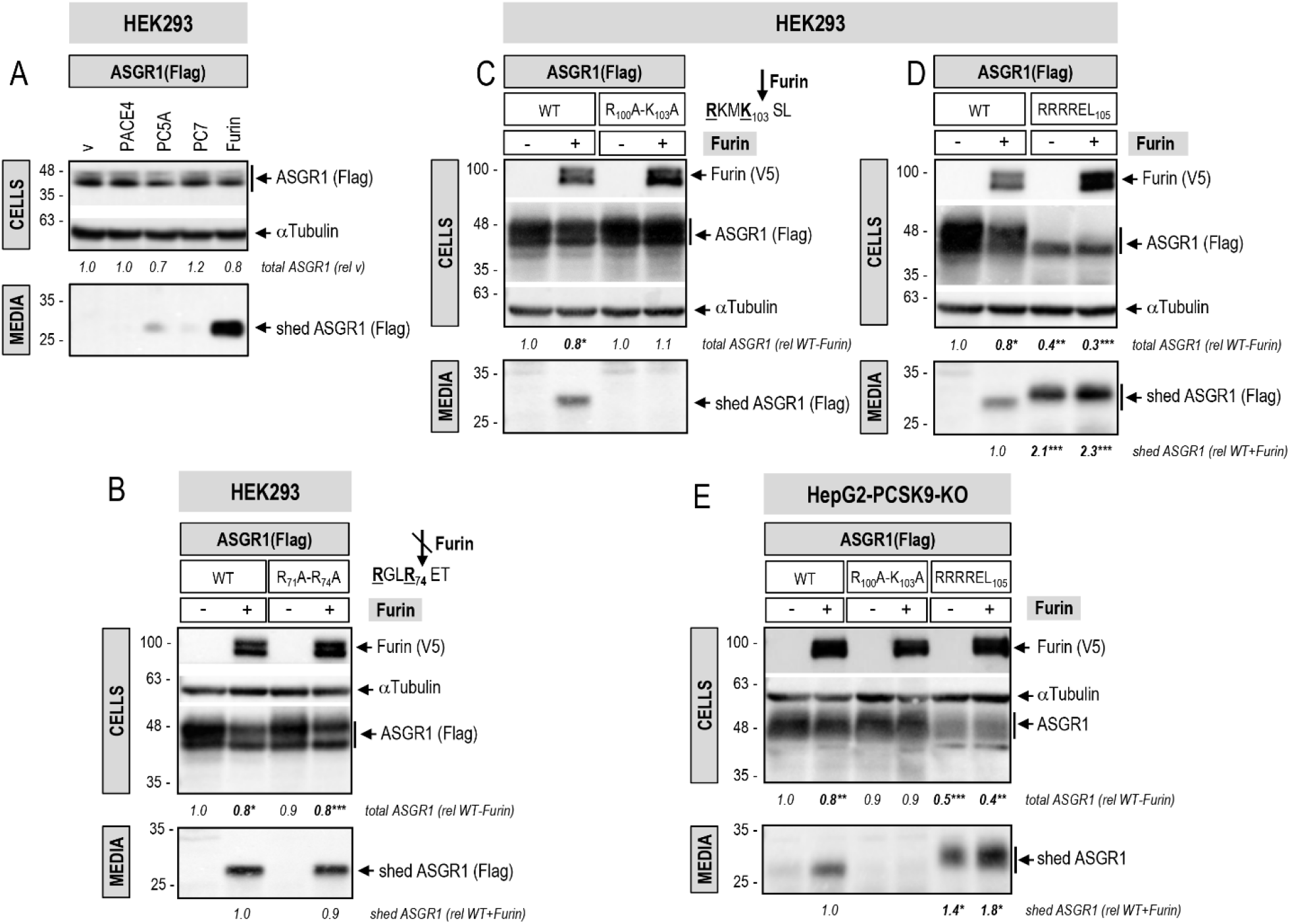
ASGR1 is shed in the media by Furin. (A) HEK293 cells were co-transfected with Flag-tagged ASGR1 and either one of the liver-expressed PCs, PACE4, PC5A, PC7, Furin, or empty vector (v), as negative control. WB analyses of cellular ASGR1 (Flag-HRP) and of shed ASGR1 (Flag-HRP) in the media are shown. HEK293 cells (B-D) or HepG2-PCSK9-KO cells (E) were co-transfected with Flag-tagged ASGR1, WT or Furin-like cleavage site mutants R_71_A-R_74_A (B), R_100_A-K_103_A (C, E), RRRREL_105_ (D, E), and V5-tagged Furin (+) or empty vector (-). WB analyses of cellular Furin (V5-HRP) and ASGR1 (Flag-HRP) and of shed ASGR1 (Flag-HRP) in the media are depicted (B-D). For HepG2-PCSK9-KO cells, ASGR1 in cells and media is revealed with ASGR1 antibody (E). The slower SDS-PAGE migration of shed ASGR1 RRRREL_105_ mutant in (D) and (E) is likely due to an increased negative charge (glutamic acid, <u>E</u>) following the Furin cleavage site in mutant protein RRRR_103_↓<u>E</u>LE compared to WT RKMK_103_↓SLE. Data are representative of three independent experiments. Quantifications are averages ± SD. * p<0.05; ** p<0.01; *** p<0.001 (t-test).

## DISCUSSION

The hepatic ASGR is pivotal for the efficient clearance of desialylated glycoproteins from circulation by receptor-mediated endocytosis and lysosomal degradation, whereupon ASGR is recycled back to the cell-surface (13,31). Glycoprotein ASGR ligands contain terminal galactose or N-acetylgalactosamine, with the latter being the preferred residues (31). In humans, the functional receptor is composed of two subunits, ASGR1 (major, ∼46 kDa) and ASGR2 (minor, ∼50 kDa) that exist as homo-oligomers or hetero-oligomers in a molar ratio of ∼3:1, respectively (13,31). Both subunits are type-II transmembrane proteins comprising a cytosolic, transmembrane domain, a luminal stalk region and a calcium-dependent CRD (Fig. 1A). Oligomerization of the ASGR subunits *via* their respective stalk regions occurs in the ER (31). ASGRs are internalized from the cell-surface *via* clathrin-coated vesicles into endosomes and are rapidly recycled constitutively even in the absence of a ligand (11). Thus, at steady-state ∼40-60% of total cellular ASGRs are on the cell-surface (31). So far, the known ligands of ASGRs are soluble desialylated proteins, including von Willebrand factor (11) and exogenous proteins (e.g. asialofuetin, asialo-orosomucoid) (32).

Recently, it was observed that conjugation of antisense oligonucleotides to N-acetylgalactosamine mediated their efficient uptake into liver hepatocytes *via* ASGR1, providing a powerful targeted delivery of these and other drugs to the liver (33). Indeed, an ASGR1-targeted antisense siRNA technology is now successfully applied to silence liver PCSK9 in hypercholesterolemic patients worldwide (34).

Circulating PCSK9 originating from hepatocytes (2) binds to cell-surface LDLR and targets it to degradation in endosomes / lysosomes (6). This mechanism led to a powerful new treatment to reduce LDLc *via* inhibitory monoclonal antibodies or liver targeted antisense siRNA silencing of its mRNA (35). In 2016, Nioi *et al*. reported that two human *ASGR1*-deletion variants found in Iceland, resulting in truncated proteins, likely inactive, are associated with a modest ∼10-14% reduction in LDLc, yet with a significant ∼34% reduced risk of coronary artery disease (12). This suggested that the atheroprotective effects of *ASGR1* LOF extended beyond reduction of serum LDLc and could implicate, extrahepatic functions of ASGR1, e.g., in macrophages (36).

Since PCSK9 is a major player in the regulation of LDLc (7) and its receptor LDLR (3,6), and ASGR1/LDLR co-localize at the cell-surface (Fig. 2A), we investigated the possible involvement of ASGR1 in the modulation of the LDLR (11) and/or PCSK9 functions. The data presented in this work clearly eliminated a reciprocal direct effect between PCSK9 and ASGR1 proteins (Figs. 2-5), and demonstrated that loss of endogenous ASGR1 in hepatic cells lines enhanced the levels of the LDLR and its ability to uptake DiI-LDL (Fig. 5), whereas the reverse is true upon overexpression of ASGR1 (Fig. 6), and both phenotypes are independent of PCSK9. This was confirmed *in vivo* in mice lacking PCSK9, which as expected exhibit higher levels of LDLR (18), but no significant change in ASGR1 protein levels (Fig. 3). Finally, lack of endogenous ASGR1 in HepG2 cells does not affect PCSK9 mRNA or its protein levels in cells and media (Fig. 4).

In the absence of PCSK9, co-IP experiments in HepG2-PCSK9-KO cells and in mouse *Pcsk9*-KO livers revealed that endogenous ASGR1 and LDLR form a complex (Figs. 8A, B). Since ASGR1 binds galactose/N-acetyl-galactosamine, and its Gln_240_, Trp_244_ and Glu_253_ are implicated in carbohydrate recognition (32), we tested the importance of these ASGR1 CRD residues in the regulation of the LDLR. The triple Ala-mutant of these residues largely failed to co-IP with mLDLR (∼150 kDa) in HEK293 cells, but still bound non-O-glycosylated iLDLR (∼110 kDa) (Fig. 8C; left panel). This unexpected result revealed that ASGR1 could bind mostly the mLDLR (N- and O-glycosylated) in a sugar-dependent fashion, and iLDLR (N-glycosylated) in an O-glycosylation-independent manner. These data suggest that the galactose/N-acetyl-galactosamine residues recognized by the CRD of ASGR1 likely reside in part on O-glycosylated chains of the mLDLR. The latter are predominantly present close to the C-terminal transmembrane domain but are also found in the linker regions separating the N-terminal repeat domains of the LDLR (22). Finally, shedding of the LDLR by metalloproteases is thought to mostly target iLDLR (23). Overexpression of WT ASGR1 decreased LDLR shedding, likely due to degradation of mainly iLDLR (Fig. 9C), and silencing endogenous ASGR1 resulted in elevated shedding of primarily iLDLR, in line with increased cellular LDLR and higher exposure to endogenous sheddase(s) (Fig. 9D). Thus, both ASGR1-induced degradation of the LDLR and the cell-surface shedding of the latter mostly target iLDLR, whereas PCSK9 targets both LDLR forms. It was of interest to note that not all cell-surface glycoproteins are targeted by ASGR1, since the endogenous IR and CD36 in HepG2 cells were insensitive to it (Fig. 10), possibly due to their low levels of desialylation.

The 9-membered family of proprotein convertases (PCs) is composed of secretory serine proteases implicated in a wide variety of functions both in health and disease (26,37). The first seven members of the family cleave substrates at the canonical motif **R/K**-Xn-**R/K**↓, where Xn=0, 2, 4 or 6 spacer aa (26). The presence of two such motifs in the primary sequence of ASGR1 suggested that it may be cleaved by one or more member(s) of the PC-family. Our data showed that only Furin can cleave ASGR1 at **<u>R</u>**KM**<u>K</u>**_103_**↓**SL and shed it into the media (Figs. 11A, C, E), but does not process the other **<u>R</u>**GL**<u>R</u>**_74_ ET site (Fig. 11B). We therefore expect that some of the hepatic ASGR1 may be circulating in plasma as a soluble shed ASGR1 form (aa 104-291) that still encodes the CRD with its critical residues (Gln_240_, Trp_244_ and Glu_253_), and conceivably could bind to non-hepatic targets and regulate their function. The generation of an optimally shed form of ASGR1 with the above sequence replaced by **<u>RRRR</u>**_103_↓**<u>E</u>**L (Figs. 11D, E) may lead in the future to the identification of some of the possible extrahepatic roles of the resulting circulating ASGR1. For example, the lack of ASGR1 in human was suggested to possibly affect receptors implicated in modulating inflammation (12).

Finally, it is interesting to note that liver ASGR1 has recently been shown to bind the N-terminal domain and ACE2 receptor binding domain of the Spike-glycoprotein of SARS-CoV-2 (38), the etiological agent of COVID-19 (39). Since liver does not significantly express ACE2, the main SARS-CoV-2 receptor (40), it is possible that ASGR1 can act as an alternative receptor in hepatocytes, thereby expanding the tropism of this deadly virus (41). It would thus be informative to define the mode of interaction of ASGR1 with the Spike-glycoprotein of SARS-CoV-2 and whether the LDLR may participate in this process.

We conclude that the LDLR is a ligand of hepatic ASGR1, representing the first case of a membrane-bound protein targeted by ASGR1 for degradation, since ASGRs are primarily known to interact with circulating factors (11,31). Our data provide a mechanism for the modest reduction of LDLc in humans lacking functional ASGR1 (12). The other functions of ASGR1 that are not related to LDLc regulation but affect inflammation, atherosclerosis and cardiovascular health (12) are yet to be identified.

## EXPERIMENTAL PROCEDURES

### Plasmids and site directed mutagenesis

The cDNAs encoding for human ASGR1 and human ASGR2 were obtained from GenScript (NJ). Human ASGR1 and its mutants (Q240A/W244A and Q240A/W244A/E253A) and human ASGR2 were sub-cloned into pcDNA 3.1 vector (Invitrogen) containing a C-terminal Flag tag (ASGR1) or HA tag (ASGR2). Point mutations of ASGR1 were created by site-directed mutagenesis, and the identity of each mutant was confirmed by DNA sequencing. The constructions containing the human LDLR, human PACE4, human PC7, human Furin or mouse PC5A, cloned in pIRES2-EGFP (Clontech), were previously described (42,43).

### Cell culture, transfections, and treatments

HepG2 (human hepatocellular carcinoma), HepG2-PCSK9-KO (CRISPR cas9 for PCSK9), IHH (immortalized human primary hepatocytes) and HEK293 (human embryonic kidney-derived epithelial) cells were cultured at 37°C under 5% CO_2_ in complete medium (Eagle’s Minimal Essential Medium for HepG2 and IHH cells; Dulbecco’s Modified Eagle’s Medium for HEK293 cells) (Gibco, Thermofisher Scientific) containing 10% (v/v) fetal bovine serum (Invitrogen).

Protein overexpression was achieved using JetPEI (PolyPlus) transfection reagent in HepG2 and HepG2-PCSK9-KO cells (2 µg total DNA/well in 12 well plate) and JetPRIME (PolyPlus) in HEK293 cells (1 µg total DNA/well in 12 well plate), following the manufacturer’s recommendations. 24 hours post-transfection, the culturing medium was changed from complete to serum-free to achieve maximal expression of the LDLR, and the cells were treated according to each experiment. Incubations with *exogenous* purified PCSK9 (ACRO Biosytems) was carried out for an additional 20 hours. LDLR functionality was assessed by incubation with DiI-LDL (Alfa Aesar, Kalen Biochemicals). Small interfering RNA (siRNA) targeted against ASGR1 or PCSK9 were purchased from siGenome, Horizon Discoveries and transfections were carried out using INTERFERin (PolyPlus) as per manufacturer’s instructions.

### Western blotting

Cells were washed twice with ice-cold PBS and lysed 60 min on ice with ice-cold non-denaturing cell lysis buffer (20 mM Tris-HCl, pH 8, 137 mM NaCl, 2 mM Na_2_EDTA, 1% Nonidet P-40, 10% glycerol, supplemented with protease inhibitor cocktail without EDTA). Mouse livers were homogenized, and protein extraction was performed in radioimmune precipitation assay buffer (50 mM Tris-HCl pH 7.8, 150 mM NaCl, 1% Nonidet P-40, 0.5% sodium deoxycholate, 0.1% SDS, supplemented with complete protease inhibitor cocktail) for 40 min on ice. Cell lysates (20-30 μg of total protein) and conditioned media (20% of total media) were electrophoretically resolved on 8 or 10% Tris-glycine SDS-polyacrylamide gels, respectively, and transferred to PVDF membranes using a Trans-Blot Turbo transfer System (Bio-Rad). Mouse liver lysates were subjected to 8% Tris-tricine SDS-PAGE before transfer to PVDF membrane. Membranes were blocked with 5% skim milk in tris-buffered saline containing Tween-20 for 1 hour and subsequently incubated with primary antibody according to the manufacturer’s recommendations. Antigen-primary antibodies complexes were detected with secondary antibodies conjugated to horse radish peroxidase and developed using a chemiluminescent reagent (Clarity ECL, Bio-Rad). Images of the WBs were acquired using a ChemiDoc MP System (Bio-Rad) and analyzed with Image Lab (version 6.0) software (Bio-Rad). Immunoblots from cell lysates and liver lysates were quantified and normalized to membranes probed for α-Tubulin or β-Actin, respectively. See Supplementary Table S1 for a list of the antibodies used for Western blot analyses. In all figures, immunoblots were cropped for clarity.

### Immunofluorescence

For immunofluorescence experiments, HEK293, HepG2 or HepG2-PCSK9-KO cells (0.5×10^5^ cells/well) were plated on poly-L-lysine-coated round microscope coverslips that were placed in a 24-well cell culture plate. Cells were then treated as required for each experiment (PCSK9 incubation, siRNA transfection, protein overexpression). To analyze plasma membrane LDLR expression, the cells were washed twice with PBS and fixed with a solution of 4% paraformaldehyde in PBS (10 min). After blocking with PBS + 2% BSA (1 hour), samples were incubated at 4 °C overnight with the respective primary antibodies (see Table S1), washed with PBS, and incubated with the appropriate fluorescent secondary antibody (see Table S1) for 1 hour at room temperature. To analyze cell-surface ASGR1, cells were incubated in SFM media with ASGR1 antibody (1/50) for 2 hours at 37°C. Cells were then fixed and incubated with appropriate secondary antibodies as described above. When LDLR functionality was tested, prior to fixation cells were incubated for 2 hours at 37°C with 5μg/ml DiI-LDL in SFM media. Coverslips were mounted on a glass slide with ProLong Gold antifade reagent with DAPI. Samples were visualized using a Plan-Apochromat 63x 1.4 oil objective of an LSM-710 confocal laser-scanning microscope (Carl Zeiss) with sequential excitation and capture image acquisition with a digital camera. Images were processed with ZEN software. Image analysis to quantify the fluorescence intensities was accomplished using Volocity®6.0.

### Animal experiments

*In situ hybridization* studies of ASGR1 and ASGR2 mRNA expression were performed on whole body tissue cryostat sections (8-to 10-μm) from WT (C57BL/6J) mice, embryonic day 10 to adult, as described previously (2,15,44). Mouse antisense and sense (negative control) cRNA (complementary RNA) riboprobes coding for ASGR1 and ASGR2 were labeled with ^35^S-UTP and ^35^S-CTP (1,250 Ci/mmol; Amersham Pharmacia), to obtain high specific activities of ≈1,000 Ci/mmol (2).

### Immunohistochemistry in mouse liver sections

LDLR and ASGR1 protein expression in mouse liver was assessed by IHC on cryosections (9-10 mice) and WB of tissue extracts (3-4 mice) from 12-16-week-old male *Pcsk9*^-/-^ and *Ldlr*^-/-^ mice on a C57BL/6J background and age-matched C57BL/6J controls. For IHC, mice liver cryosections (8 µm thick) were fixed in 4% paraformaldehyde in PBS for 1 hour, rinsed in PBS with 0.1% glycine, washed in PBS, and blocked in 1% BSA (Sigma-Aldrich, Oakville, ON) in PBS for 1 hour at room temperature. Sections were then incubated overnight at 4°C with the LDLR and ASGR1 primary antibodies (see Table S1) and washed three times for a total of 15 min in PBS. Labeling was visualized by incubation with Alexa Fluor 488-labeled secondary antibodies (see Table S1) for 1 hour at room temperature in PBS. After three washes with PBS, nuclei were counterstained with Hoechst dye (Sigma-Aldrich, Oakville, ON). Images were acquired as described previously using a LSM 700 confocal microscope equipped with ZEN 2011 software (45). Data quantification was achieved using Matlab software. For all images the minimum threshold was set to a value of 20, negative control values were subtracted, and final values were analyzed using Microsoft Excel. For each genotype, the most representative image was chosen.

### Quantitative RT-PCR

Quantitative RT-PCR was performed as previously described (18). Briefly, total RNA from liver or cells was extracted with TRIzol (Invitrogen). cDNA was generated from 1µg of total RNA using a SuperScript IV cDNA reverse transcriptase (Invitrogen). Quantitative PCR was performed using the SYBR Select Master Mix (Applied Biosystems, Carlsbad, CA) and the ΔΔct method. Expression of each human gene was normalized to that of Tata-binding protein (TBP), while expression of the mouse genes was normalized to that of hypoxanthine phosphoribosyl transferase (HPRT). The sets of primers (ThermoFisher Scientific) were as follows: human ASGR1, 5’
s-GGGAAGAAAGATGAAGTCGCTAGA *versus* 5’-GCAGGCTGGAGTGATCTTCAC; human ASGR2, 5’-GACGGAGGTCCAGGCAATC *versus* 5’-TGGCTCCTAGGGATGTGATCTT; human LDLR, 5’-AGGAGACGTGCTTGTCTGTC *versus* 5’-CTGAGCCGTTGTCGCAGT; human PCSK9, 5’-TGGAGCTGGCCTTGAAGTTG *versus* 5’-GATGCTCTGGGCAAAGACAGA; human SREPB2, 5’-AGAATGTCCTTCTGATGTCC *versus* 5’-GGAGAGTCTGGCTCATCTT; human TBP, 5’-CGAATATAATCCCAAGCGGTTT *versus* 5’-GTGGTTCGTGGCTCTCTTATCC; mouse ASGR1, 5’-TCTGACGTGCGAAGCTTGAG *versus* 5’-GGTCCTTTCAGAGCCATTGC; mouse ASGR, 5’-CGATGATGAACATGGCTCTCA *versus* 5’-AGGCTGCCCTTTCCAGTGT; mouse HPRT, 5’-CCGAGGATTTGGAAAAAGTGTT *versus* 5’-CCTTCATGACATCTCGAGCAAGT.

### Co-immunoprecipitation

For co-immunoprecipitation of LDLR-ASGR1 complexes from HepG2-PCSK9-KO cells (endogenous LDLR-ASGR1 complex) and transfected HEK293 cells, cells were lysed in Pierce non-denaturing IP buffer supplemented with complete protease inhibitor cocktail. Lysates containing 0.5 mg total protein (HepG2-PCSK9-KO cells) or 0.25 mg total protein (HEK293 cells) were exposed overnight, at 4 °C on a rocker, to 2 µg of IP antibody (rabbit anti-ASGR1 antibody for HepG2-PCSK9-KO cell lysates; mouse (IgG1)MAB Flag M2 or mouse (IgG2a) anti-V5 MAB antibodies for HEK293 cell lysates). The following day, 60 µl True Blot anti-rabbit IgG beads (HepG2-PCSK9-KO cell lysates) or 40 µl of protein A/G PLUS-Agarose (HEK293 cell lysates) were added for an additional 1 to 2 hours incubation. Following three washes with lysis buffer/protease inhibitors and two washes with PBS and elution in 70 µl 2x Laemmli sample buffer, the pull-downs were separated by 8% Tris-glycine SDS-PAGE along with inputs (10% of original material used for co-IP) and analyzed by WB for LDLR and ASGR1 using their respective primary antibodies. In the case of LDLR-ASGR1 co-immunoprecipitation from HepG2-PCSK9-KO cell lysates, separate pull-downs were also analyzed for endogenous CD36 and IR as negative controls for membrane-bound receptors. Rabbit IgG TrueBlot was used for the detection of immunoblotted ASGR1 from HepG2-PCSK9-KO cell lysates (endogenous LDLR-ASGR1 complex), which eliminated the hindrance by interfering immunoprecipitating immunoglobulin heavy and light chains. TrueBlot preferentially detects the non-reduced form of rabbit IgG over the reduced, SDS-denatured form of IgG.

For co-immunoprecipitation of LDLR-ASGR1 complexes from mice livers, fresh livers from *Pcsk9*-KO mice littermates were lysed in Pierce non-denaturing IP buffer supplemented with complete protease inhibitor mixture. Liver lysates containing 1 mg of total protein were exposed to 2 µg of rabbit anti-ASGR1 antibody overnight at 4 °C on a rocker. The following morning, 100 µl of Surebeads Protein G magnetic beads (BioRad) was added to each sample tube, including “beads only” negative control. Isolation and purification was carried out using magnets as per manufacturer’s instructions. A “control” was included, which consisted of a liver lysate from a wild type (C57BL/6J) mouse to which ASGR1 antibody was added to help identify the IgGs. See Table S1 for antibodies used.

### Cell-surface biotinylation

Biochemical detection of cell-surface LDLR and ASGR1 by Western blot analysis was performed following the protocol described in (46). Namely, HEK293 cells seeded in 6-well plates and after reaching 80% confluency were transiently co-transfected with 1.5 µg DNA consisting of V5-tagged LDLR and empty vector or a combination of LDLR-V5, HA-tagged ASGR2 and Flag-tagged ASGR1, WT or carbohydrate-binding mutants Q240A/W244A/E253A or Q240A/W244A.

Following 48 hours post-transfection, cells were washed twice with ice-cold PBS and biotinylated with 2 ml of EZ-link sulfo-NHS-LC-Biotin (ThermoScientific 0021335, ON) (0.5 mg/ml in PBS) for 20 min at 4°C, on ice. After 10 min incubation on ice with ice-cold BSA (0.1 mg/ml) in PBS to quench the reaction and two washes with cold PBS, cells were lysed for 10 min on ice in biotinylation lysis buffer (20 mM Tris pH 7.5, 5 mM EDTA, 5 mM EGTA, 0.5% N-dodecyl-N-maltoside, supplemented with protease inhibitor cocktail without EDTA). Cell lysates were collected after centrifugation at 3,000 rpm for 10 min at 4°C and quantified for protein concentration. Biotinylated cell-surface proteins (240 µg total protein) were immunoprecipitated overnight with 30 µl Streptavidin agarose. Following centrifugation at 7,000 rpm for 2 min at 4°C, the pellet containing the plasma membrane pool was washed three times, eluted in 60 µl 2x Laemmli sample buffer and separated by 8% Tris-glycine SDS-PAGE. A negative control (-Biotin) was included, to which PBS was added instead of Biotin at the biotinylation step. Total proteins (plasma membrane + intracellular) were detected from a fraction of the lysates before Streptavidin immunoprecipitation, which was referred to as “input”.

### Statistical analysis and graphical representation

Quantifications are defined as averages ± S.D. The statistical significance was evaluated by Student’s *t* test, and probability values (*p*) <0.05 were considered significant. Data representation was performed using GraphPad Prism 8 software.

## AUTHORS CONTRIBUTIONS

NGS, DSR, EG and RCA conceived the studies and wrote the manuscript. DSR, EG, RE, AR, JM, RMD, AE, JHB and PFL performed all experiments.

## SOURCES OF FUNDING

This work was supported in part by research grants to Nabil G. Seidah from the CIHR Foundation Scheme grant (148363), a Canada Research Chair (950-231335) and a Leducq Foundation grant (13 CVD 03), as well as to Richard C. Austin from the Heart and Stroke Foundation of Canada (G-13-0003064, G-15-0009389) and the Canadian Institutes of Health Research (74477, 173520). Financial support from the Research Institute of St. Joe’s Hamilton is acknowledged. Richard C. Austin is a Career Investigator of the Heart and Stroke Foundation of Ontario and holds the Amgen Canada Research Chair in the Division of Nephrology at St. Joseph’s Healthcare and McMaster University.

## CONFLICT OF INTEREST

The authors declare that they do not have anything to disclose regarding funding or conflict of interest regarding this manuscript.

## Notes

### Competing Interest Statement

The authors have declared no competing interest.

### Summary of Updates

This revised version is an updated version with some new data on the immunocytochemistry of ASGR1

