## supplemental table S1 and Figure S1 for "Asialoglycoprotein receptor 1 is a novel PCSK9-independent ligand of liver LDLR that is shed by Furin"

**Supplementary Table S1. Antibodies used for Western blotting, immunofluorescent staining, immunohistochemical staining and immunoprecipitation**

| Antibody | Catalog No; Source | Application | Dilution |
|---|---|---|---|
| **human LDLR** | AF2148; R&D | WB | 1:1,000 |
| **mouse LDLR** | AF2255; R&D | WB | 1:1,000 |
| **ASGR1** | 11739-1-AP; Proteintech | WB | 1:1,000 |
| **α-Tubulin** | 11224-1-AP; Proteintech | WB | 1:5,000 |
| **μ-Actin** | A2066; Sigma | WB | 1:5,000 |
| **Insulin receptor β** | ENZ-ABS710; Enzo | WB | 1:1,000 |
| **CD36** | Ab36977; Abcam | WB | 1:1,000 |
| **human PCSK9** | (1) | WB | 1:3,000 |
| **Flag-HRP** | A8592; Sigma | WB | 1:5,000 |
| **HA-HRP** | H6533; Sigma | WB | 1:4,000 |
| **V5-HRP** | V2260; Sigma | WB | 1:10,000 |
| **Rabbit TrueBlot®: Anti-Rabbit IgG HRP** | 18-8816-31; Rockland Inc. | WB | 1:2,000 |
| **human LDLR** | AF2148; R&D | IF | 1:200 |
| **ASGR1** | 11739-1-AP; Proteintech | IF | 1:200 |
| **mouse LDLR** | AF2255; R&D | IHC | 1:150 |
| **ASGR1** | 11739-1-AP; Proteintech | IHC | 1:300 |
| **Goat anti rabbit IgG Alexa Fluor 488** | A27034; Invitrogen | IHC/IF | 1:150/1:500 |
| **Donkey anti goat IgG Alexa Fluor 488** | A-11055; Invitrogen | IHC/IF | 1:150/1:500 |
| **Flag M2** | F1804; Sigma | IP/IF | n/a /1:250 |
| **V5** | R960-25; Invitrogen | IP | n/a |
| **ASGR1** | 11739-1-AP; Proteintech | IP | n/a |

WB, Western blot; IF, immunofluorescence; IHC, immunohistochemistry

1. Nassoury, N., Blasiole, D. A., Tebon, O. A., Benjannet, S., Hamelin, J., Poupon, V., McPherson, P. S., Attie, A. D., Prat, A., and Seidah, N. G. (2007) The Cellular Trafficking of the Secretory Proprotein Convertase PCSK9 and Its Dependence on the LDLR. *Traffic.* **8**, 718-732

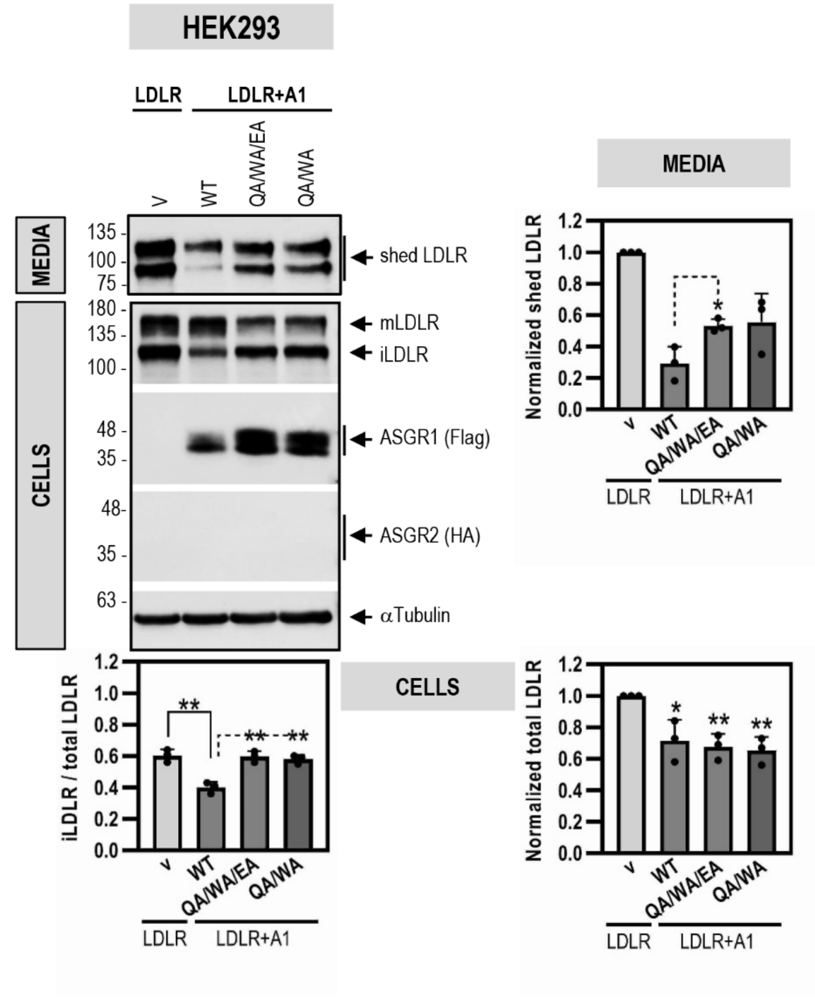


**Supplementary Figure S1: The ASGR1-mediated degradation of LDLR through its lectin-binding domain is independent of ASGR2 when co-expressed in HEK293 cells**

HEK293 cells were co-transfected with V5-tagged LDLR and either Flag-tagged ASGR1 (A1), WT or its carbohydrate-binding mutants Q240A/W244A/E253A (QA/WA/EA) or Q240A/W244A (QA/WA), or control empty vector (v). WB analyses of total cellular LDLR (mature mLDLR, and immature iLDLR) and ASGR1 (Flag-HRP) and of shed LDLR in the media are shown. Quantifications are averages ± SD of three independent experiments. * $p<0.1$; ** $p<0.01$ (t-test) are relative to the control vector condition.
